# LIMD1 Loss Confers a GPX4-Dependent Cell State in Lung Cancer

**DOI:** 10.1101/2025.06.06.658274

**Authors:** Akash Saha, Paul Grevitt, Jair Marques Junior, Alex Von Kriegsheim, Kunal Shah, Zhen Gao, Brent R. Stockwell, Andrew Finch, Barrie Peck, Tyson V. Sharp

**Author notes:** Co-corresponding Author: Paul Grevitt. Corresponding Author: Prof Tyson V. Sharp.

## Abstract

*LIMD1* (LIM domains containing 1) is a bona-fide tumour suppressor gene frequently lost during the early stages of non-small cell lung cancer (NSCLC) development, resulting in a worse outcome for *LIMD1*-deficient patients. *LIMD1* deficiency is present in approximately 50% of NSCLC cases, representing at least 21,000 patients in the United Kingdom and 1.2 million worldwide. During cancer progression, cells can accumulate genetic changes that render them more dependent on certain genes for their survival. By performing CRISPR-Cas9 dropout screens, we identified *GPX4* as a novel vulnerability in *LIMD1* deficient LUAD cells. By targeting *GPX4* using RNA interference (RNAi) and pharmacological intervention in our isogenic LUAD lines along with a non-transformed lung cell line, we validated GPX4 as a novel dependency in *LIMD1* deficient cells. GPX4 is a key defence mechanism against ferroptosis, an iron-dependent form of regulated cell death. This state of increased ferroptosis susceptibility upon *LIMD1* loss is due to increased basal reactive oxygen species levels and lipid peroxidation. Importantly, targeting GPX4 with ferroptosis inducing agents in NSCLC patients with *LIMD1* loss may represent a novel therapeutic strategy.

## INTRODUCTION

Lung cancer (LC) remains the leading cause of cancer-related morbidity and mortality globally, with approximately 2 million new cases and 1.8 million deaths each year ^1^. In the UK, LC is the third most common cause of cancer-related death, accounting for 21% of all cancer fatalities and over 34,800 deaths annually ^2^. It also ranks as the third most frequently diagnosed cancer in the UK, with more than 48,500 new cases annually ^2^. LC is broadly classified into small-cell lung carcinoma (SCLC) and non-small cell lung carcinoma (NSCLC), constituting 15% and 85% of cases, respectively ^3^. NSCLC is further subdivided into large-cell carcinoma, lung adenocarcinoma (LUAD), and lung squamous cell carcinoma (LUSC), with LUAD and LUSC representing 40% and 25-30% of all LC cases ^4^.

Disease stage at diagnosis is the most critical determinant of patient prognosis and therapeutic options ^5^. Early-stage NSCLC (stages I-II) is typically treated with surgical resection, offering 5-year survival rates of 64.6% and 41.2%, respectively ^6^. In contrast, advanced-stage NSCLC (stages III-IV) has significantly poorer 5-year survival outcomes of 36-13% and 1%, respectively ^7,8^. Treatment for advanced disease often includes chemotherapy and radiation, but outcomes remain dismal ^1^. A key challenge in LC management is the high frequency of late diagnosis due to symptom overlap with other chronic respiratory diseases, resulting in more than 60% of patients presenting with advanced-stage diseases ^9^. Despite recent advances in immunotherapy and tyrosine kinase inhibitors (TKIs), overall 5-year survival rates remain stagnant at 18% ^10,11^, underscoring the urgent need for novel, personalised treatment approaches.

LIM domain-containing protein 1 gene (LIMD1), a 73 kDa tumour suppressor encoded within the common eliminated region 1 (C3CER1) on chromosome 3p21.3, plays a critical role in LC tumorigenesis ^12,13^. Deletions in this region are frequent across multiple cancers, including breast, colorectal ovarian, renal, and gastric malignancies ^14^. LIMD1 is subject to both genomic deletion and epigenetic silencing via promoter hypermethylation, which represses transcriptional activity ^12,15^. Importantly, LIMD1 loss occurs early in NSCLC development: approximately 46% of LUAD and 85% of LUSC patients exhibit monoallelic deletion, with TRACERx data indicating this loss is often clonal ^16,17^. Copy number correlates strongly with mRNA expression levels in both LUAD and LUSC, and patients with LIMD1 loss show reduced expression ^16^. TCGA data further confirm significantly lower LIMD1 mRNA expression in tumour samples compared to adjacent normal tissue, which is associated with worse prognosis in LUAD ^16^.

It has been robustly demonstrated that LIMD1 loss drives NSCLC progression ^*18*^. LIMD1 overexpression in LUAD cell lines suppresses colony formation, indicating a growth-suppressive role ^18^. *In vivo, Limd1*^*-/-*^ mice develop significantly more and larger lung tumours upon urethane exposure or in the *KRAS*^*G12D*^ background ^12^.

LIMD1 is a member of the Zyxin family of LIM domain containing proteins, characterised by an unstructured N-terminal pre-LIM domain and three tandem zinc-finger LIM domains at the C-terminus, facilitating protein-protein interactions ^19^. LIM domains are evolutionarily conserved modules that regulate diverse processes, including gene expression, signalling, and cell adhesion ^19^. As a multifunctional scaffold, LIMD1 integrates signals across multiple homeostatic pathways ^20^. Notably, LIMD1 binds the retinoblastoma protein (pRb), suppressing E2F1-driven transcription and, cell cycle progression ^18^. It also modulates the hypoxic response by promoting efficient hydroxylation and proteasomal degradation of HIF -1α via scaffolding and recruiting a PHD-pVHL complex ^16,21^. More recently, LIMD1 was shown to be important for microRNA-induced silencing complex (miRISC) function through site-specific phosphorylation ^22,23^.

Given its pleiotropic regulatory roles, LIMD1 loss disrupts multiple tumour-suppressive pathways, promoting tumour initiation, progression, and metastasis ^12,16,18,21^. However, the multi-faceted nature of LIMD1 biology complicates direct therapeutic targeting. A prior drug repurposing screen identified PF-477736 as selectively cytotoxic to *LIMD1*-deficient NSCLC cells ^20^. Although originally designed as a Chk1 inhibitor, its mechanism of action was independent of Chk1, functioning instead as a broad-spectrum kinase inhibitor ^20^. Despite showing *in vitro* and *in vivo* efficacy, the undefined mechanism limits clinical translation, highlighting the need for better LIMD1 targeted strategies.

Given the early and frequent loss of *LIMD1* in NSCLC, synthetic lethality offers a promising route to selectively target *LIMD1*-deficient tumours. Synthetic lethality arises when simultaneous disruption of two genes leads to cell death, whereas loss of either alone is tolerated ^24,25^. This approach enables targeted elimination of tumour cells harbouring specific vulnerabilities, sparing normal cells.

To identify such vulnerabilities, we conducted genome-wide CRISPR-Cas9 dropout screens in isogenic *LIMD1*^*+/+*^ and *LIMD1*^*-/-*^cells. This revealed a dependency in *LIMD1*-deficient cells on components of the selenocysteine biosynthesis and incorporation pathway – including EEFSEC, SEPSECS, and PSTK – and critically, on the selenoprotein GPX4. GPX4 is a key antioxidant enzyme that prevents ferroptosis, an iron-dependent form of regulated cell death triggered by uncontrolled lipid peroxidation in polyunsaturated phospholipids ^26-28^.

Here, we functionally validate GPX4 as a synthetic lethal target in *LIMD1*-deficient cells using both siRNA-mediated knockdown and pharmacological inhibition. Our results across LUAD cell lines and isogenic small airway epithelial cells (SAECs) demonstrate a clear and selective vulnerability, positioning GPX4 as a promising therapeutic target for a genomically defined NSCLC subset.

## MATERIALS AND METHODS

### Cell lines and culture conditions

A549 cells were cultured in Dulbecco’s Modified Eagle’s Medium (DMEM; Gibco), while H1299 cells were maintained in Roswell Park Memorial Institute 1640 Medium (RPMI 1640; Gibco) or Plasmax (Ximbio, #156371). SAECs were cultured using Promocell Small Airway Epithelial Cell Basal Medium (PromoCell; C-21270), supplemented with the provided Supplement mix (PromoCell; C-39170). All culture media were supplemented with 10% (v/v) fetal bovine serum (FBS; Sigma-Aldrich) and 1% (v/v) penicillin-streptomycin (ThermoFisher Scientific). Cells were incubated at 37 °C, with 5% CO_2_ and passaged at 80% confluency. Mycoplasma testing was routinely performed, and cells were used up to passage 10. Cell numbers were quantified using a Countess II automated cell counter (ThermoFisher Scientific) to ensure optimal seeding density for each assay.

### CRISPR-Cas9 dropout screen

HEK293T cells were transfected with Cas9 endonuclease to generate viral particles for subsequent transduction of isogenic *LIMD1*^+/+^ and *LIMD1*^*-/-*^ A549 cells. The Human Improved Genome-Wide Knockout CRISPR Library v1 (Addgene #67989), containing 90,709 guide sequences targeting a total of 18,010 genes, was cloned into a lentiviral plasmic backbone ^29^. *LIMD1*^+/+^ and *LIMD1*^*-/-*^ cells expressing Cas9 were transduced at low multiplicity of infection to ensure each cell received a single sgRNA. Screens were performed in two biological replicates, and cells were harvested at early and late timepoints for negative selection analysis ^30^. The relative abundance of each sgRNA was compared between genotypes to identify differential dependencies.

### siRNA transfection

The following siRNAs were purchased from Merck/Sigma-Aldrich: GPX4 (SASI_Hs02_00308687), GPX4_2 (SASI_Hs01_00050315), PLK1 (SASI_Hs01_00194343), ACSL4 (SASI_Hs01_00114667), UBB (SASI_Hs01_00201423), and a non-targeting control (#SIC001). Additionally, ON-TARGETplus SMARTpool siRNAs targeting GPX4 (L-011676-00-0005) and corresponding non-targeting controls (D-001810-10-05) were obtained from Dharmacon reagents. siRNAs were transfected using DharmaFECT 1 (Horizon Discovery) and Opti-MEM (ThermoFisher) according to manufacturer’s protocol.

### Western blotting

Cells were lysed in RIPA buffer (50 mM Tris-HCl [pH 8.0], 150 mM NaCl, 1% NP-40, 0.5% sodium deoxycholate, 0.1% SDS, supplemented with protease inhibitors (Roche, #04906837001) and phosphatase inhibitors (ThermoFisher, #A32955). Protein concentration was quantified using the BCA assay (ThermoFisher, #23225). Samples (20-30 µg) were prepared in 5x SDS loading buffer, boiled at 95 °C for 5 minutes, and resolved on 12% Bis-Tris or Tris-Glycine gels. Proteins were transferred onto PVDF membranes (Sigma, #IPVH00010), blocked, and probed with the following primary antibodies: LIMD1 (1:2000; in-house), LIMD1 (1:1000; Cell Signaling #13245), GPX4 (1:1000; Cell Signaling #52455S), and β-actin (1:20000 dilution; Sigma #A5441). HRP-conjugated secondary antibodies (Agilent) were used for detection via enhanced chemiluminescence (ThermoFisher #32106; Merck #WBKLS0500). Densitometry was performed using ImageJ 1.53e (NIH), and protein levels were normalised to β-actin or vinculin, from the same membrane.

### Cell viability assay (CellTiter-Glo)

Cells were seeded in 96-well plates and either transfected with siRNA or treated 24 hours with a dose range of test compounds or vehicle. A549 cells received two treatments, RSL3 or erastin, 48 hours apart. H1299 and SAECs received a single 24-hour treatment. At endpoint, cell viability was measured using the CellTiter-Glo 2.0 assay (Promega, #G9241), diluted 1:4 with PBS. 100 µL of reagent was added per well, plates were shaken for 10 minutes at room temperature, and luminescence was measured after transferring 80 µL were transferred to white 96-well plates.

### Crystal violet colony formation assay

Cells were seeded in 6-well plates and treated 24 hours later with drugs or siRNA (reverse transfection). Media were refreshed every 2 – 3 days. Ten days post-treatment, cells were fixed with methanol and stained with 0.05% crystal violet. Colony intensity was quantified using ImageJ.

### IncuCyte live cell imaging

Cells were seeded into 96-well plates at an optimised density and either reverse-transfected with siRNA or treated with drug/vehicle 24 hours later. Plates were imaged every 3 hours using the IncuCyte ZOOM system (Essen Bioscience) at 10x magnification. Confluence was quantified using IncuCyte ZOOM software.

### Flow cytometry

Cells (1.25×10^5^ cells per well) were seeded in 6-well plates, treated for 24 hours with drug or vehicle, and then stained for ROS and lipid peroxidation. Positive controls were treated with 300 µM H_2_O_2_ (lipid peroxidation) or 100 µM TBHP (ROS) for 15 – 20 minutes. Cells were trypsinised, neutralised with 2% FBS in PBS (FACS buffer), pelleted, and resuspended in 500 µL FACS buffer containing either 2 µM BODIPY 581/591 C11 (ThermoFisher #D3861) or 100 µM H2DCFDA (Tocris #5935). Cells were incubated for 15 minutes at 37 °C in the dark, washed twice, and passed through a 35 µm strainer polystyrene tubes (Corning #352235). A minimum of 10,000 events were collected per sample using a BD LSR Fortessa cytometer (FITC 488 nm and PE 582 nm lasers), and data were analysed in FlowJo v10 using appropriate gating.

### Oil Red O staining

Isogenic H1299 cells were seeded onto coverslips in 6-well plates (1.25 × 10^5^ cells per well). After 24 hours, cells were treated with 100 µM of BSA-oleate (Cayman #29557) or BSA control (Cayman #29556) for another 24 hours. Oil Red O working solution was prepared by mixing 6 mL of 5 mg/mL stock with 4 mL distilled water, incubating for 10 minutes, and filtering. Cells were fixed in 4% PFA for 15 minutes on ice, washed in PBS, incubated with 60% isopropanol for 5 minutes, then stained with Oil Red O for 15 minutes. After rinsing, cells were counterstained with haematoxylin (Vector H-3502), followed by a bluing agent. Coverslips were mounted using Vectashield (Vector H-1000) and imaged on a Zeiss microscope.

### Statistical analysis

GraphPad Prism, v9.4.0 was used for all statistical analyses. Data were normalised to relevant controls. Appropriate tests were selected based on group number and data. Dunnett’s multiple comparisons test was used for comparisons against a single control. Significance thresholds were defined as follows: *p < 0.05, **p < 0.01, ***p < 0.005, ****p < 0.0001.

## RESULTS

### CRISPR-Cas9 screens identify GPX4 as a synthetic vulnerability in *LIMD1*-deficient lung cancer cells

To uncover novel synthetic dependencies arising from *LIMD1* loss, we performed whole-genome CRISPR-Cas9 dropout screens in isogenic A549 LUAD cells (*LIMD1*^*+/+*^ and *LIMD1*^*-/-*^), using a pooled sgRNA library targeting 18,010 human genes (90,709 guides; Fig. 1A), with cells harvested at early and late timepoints. Analysis of sgRNA abundance revealed distinct vulnerabilities between genotypes: *LIMD1*-proficient cells were enriched for dependencies in oxidative phosphorylation and the TCA cycle, including NDUFA1, NDUFA6, NDUFA8, NDUFC2, NDUFV1, ACAD9, FOXRED1, UQCR10, COA7, COX18, and SCO2 (Fig. 1B, C). In contrast, *LIMD1*-deficient cells displayed significant enrichment for genes involved in selenocysteine (Sec) tRNA biosynthesis and incorporation pathway, including EEFSEC, SEPSECS, and PSTK. Notably, the sole Sec-containing enzyme among the enriched hits was glutathione peroxidase 4 (GPX4) – a selenoprotein that suppresses ferroptosis by detoxifying lipid hydroperoxides in cellular membranes ^26,31^.

**Figure 1.**
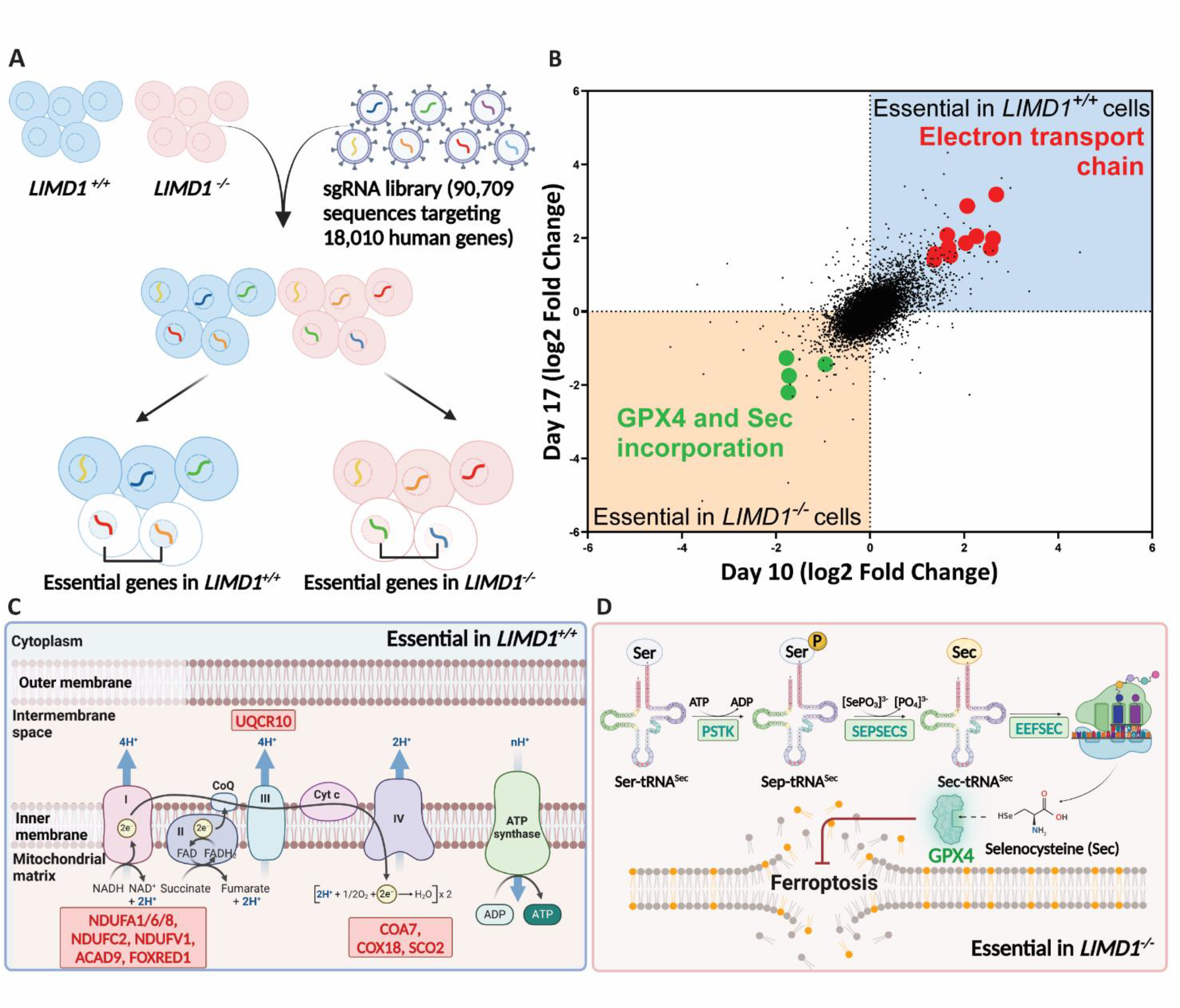
CRISPR-Cas9 dropout screens reveal dependency on selenoprotein GPX4 and enzymes involved in Sec-tRNA biosynthesis and incorporation. **(A)** CRISPR-Cas9 dropout screens performed in CRISPR-Cas9 generated isogenic A549 cells using a sgRNA library of 90,709 sequences targeting 18,010 human genes with a coverage of 5 sgRNAs per gene. **(B)** Dot-plot showing essential genes in *LIMD1*^*-/-*^ (green) and *LIMD1*^*+/+*^ cells (red) upon sgRNA abundance analysis following screen performed at an early time-point (10 days) and late time-point (17 days). **(C-D)** Schematic diagram revealing enzymes involved in ETC as essential vulnerabilities in *LIMD1*^*+/+*^ **(C)** whereas enzymes involved in Sec t-RNA biosynthesis and incorporation along with GPX4 are essential in *LIMD1*^*-/-*^ A549 cells **(D)**.

Ferroptosis, first described by Dixon et al. in 2012, is an iron-dependent form of regulated cell death driven by lipid peroxidation of polyunsaturated fatty acid (PUFA)-containing phospholipids ^26,32^. GPX4 is a key ferroptosis defence enzyme and has emerged as a therapeutic target in multiple cancers, including NSCLC ^*32,33*^.

### *GPX4* silencing via RNAi selectively impairs viability of *LIMD1*-deficient LUAD and SAEC cells

To validate *GPX4* dependency, we used siRNAs targeting *GPX4* in A549 isogenic clones. Colony formation assays revealed that *LIMD1*^-/-^ clones (C5, C10) exhibited significantly impaired clonogenic potential compared to the *LIMD1*^*+/+*^ (C3) upon *GPX4* knockdown (Fig. 2A–C). Similar results were obtained in H1299 LUAD isogenic lines using two independent siRNA sequences. Live cell imaging showed that both *LIMD1*^-/-^ clones (C25, C59) exhibited decreased confluence upon *GPX4* knockdown compared to *LIMD1*^*+/+*^ clones (C8, C9), while PLK1 siRNA served as a positive control for cytotoxicity (Fig. 2D, E; Fig. S1A, B). Cell viability assays confirmed this genotype-selective effect (Fig. S1C).

**Figure 2.**
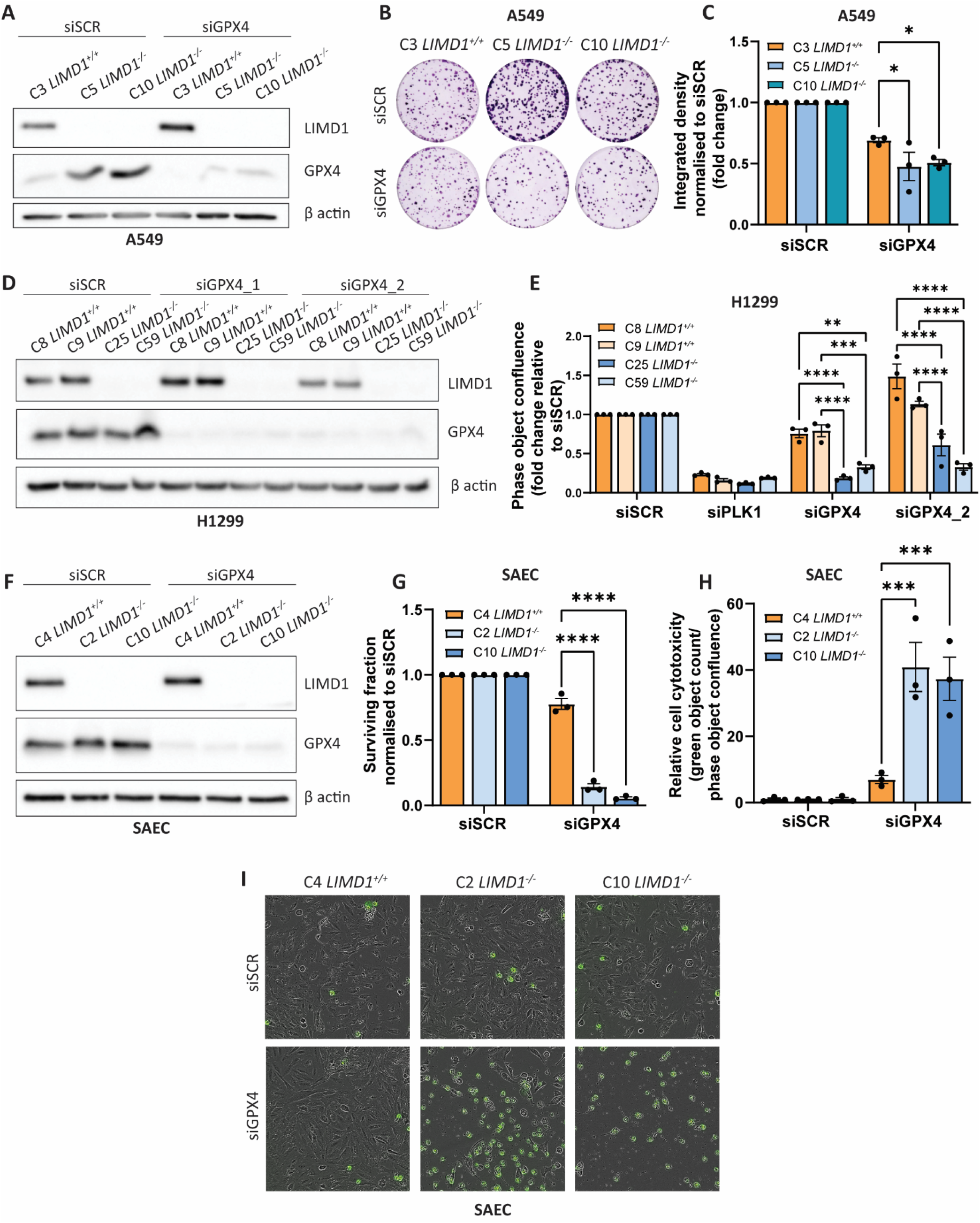
RNA interference-mediated *GPX4* targeting selectively inhibits *LIMD1*-deficient cells. **(A)** Immunoblot of isogenic A549 cells upon upon 30 nM siRNA treatment for 48 h. **(B)** Representative images from colony formation assay of isogenic A549 cells, seeded at 500 cells per well, upon treatment with 30 nM siRNA for 10 days, with growth medium being replenished every 2-3 days. **(C)** Integrated density quantification of colony formation assay images, shown as fold-change values normalised to the siSCR group for each cell line (n=3). **(D)** Immunoblot of isogenic H1299 cells upon upon 30 nM siRNA treatment for 48 h. **(E)** Phase object confluence levels of isogenic H1299 cells, seeded at 500 cells per well, upon 83 h reverse-transfection with 30 nM siRNA (n=3). **(F)** Immunoblot of isogenic SAEC cells upon upon 30 nM siRNA treatment for 48 h. **(G)** Surviving fractions of isogenic SAEC cells, seeded at 2000 cells per well, upon 96 h reverse-transfection with 30 nM siRNA (n=3). **(H, I)** Cell cytotoxicity levels **(H)** and associated-representative images **(I)** of isogenic SAEC cells, seeded at 2000 cells per well, upon 90 h reverse transfection with 30 nM siRNA (n=3). Data is shown as biological replicates ± S.E.M. Two-way ANOVA with Dunnett’s multiple comparisons test was performed in **C, G** and **H**, whereas Two-way ANOVA with Tukey’s comparisons test was performed in **E**. \**p* ≤ 0.05, \*\**p* ≤ 0.01, \*\*\**p* ≤ 0.001, \*\*\*\**p* ≤ 0.0001.

To extend these findings to a non-transformed background to exclude cooperation between *LIMD1* loss and other oncogenic drivers, we used isogenic primary human SAECs. SMARTpool siRNA-mediated *GPX4* knockdown in *LIMD1*^-/-^ SAECs (C2, C10) significantly decreased cell viability compared to *LIMD1*^*+/+*^ cells (C4), with ferrostatin-1 (Fer-1) rescuing this effect, confirming ferroptosis as the mode of cell death (Fig. 2F, G; Fig. S1D). Live cell imaging similarly revealed selective growth inhibition in *LIMD1*^-/-^ SAECs (Fig. S2E, F). Finally, cell membrane integrity assays confirmed cytotoxic (not merely cytostatic) effects of *GPX4* silencing in *LIMD1*^-/-^ SAECs (Fig. 2H, I; Fig. S2g). These results demonstrate that *LIMD1*-deficient LUAD and normal airway cells are selectively dependent on *GPX4* for survival, consistent with a synthetic lethal interaction.

### GPX4 inhibition using ferroptosis-inducing agents confirms enhanced sensitivity in *LIMD1*-deficient cells

To pharmacologically validate the dependency on *GPX4*, we treated isogenic A549, H1299, and SAECs with multiple ferroptosis inducing agents (FINs), including class I (erastin, IKE), class II (RSL3, ML210), and class III (FIN56) inhibitors ^26,34-36^. In A549 isogenic cells, RSL3 and erastin treatment selectively impaired viability of *LIMD1*^-/-^ cells, which was rescued by Fer-1 co-treatment (Fig. 3A–D).

**Figure 3.**
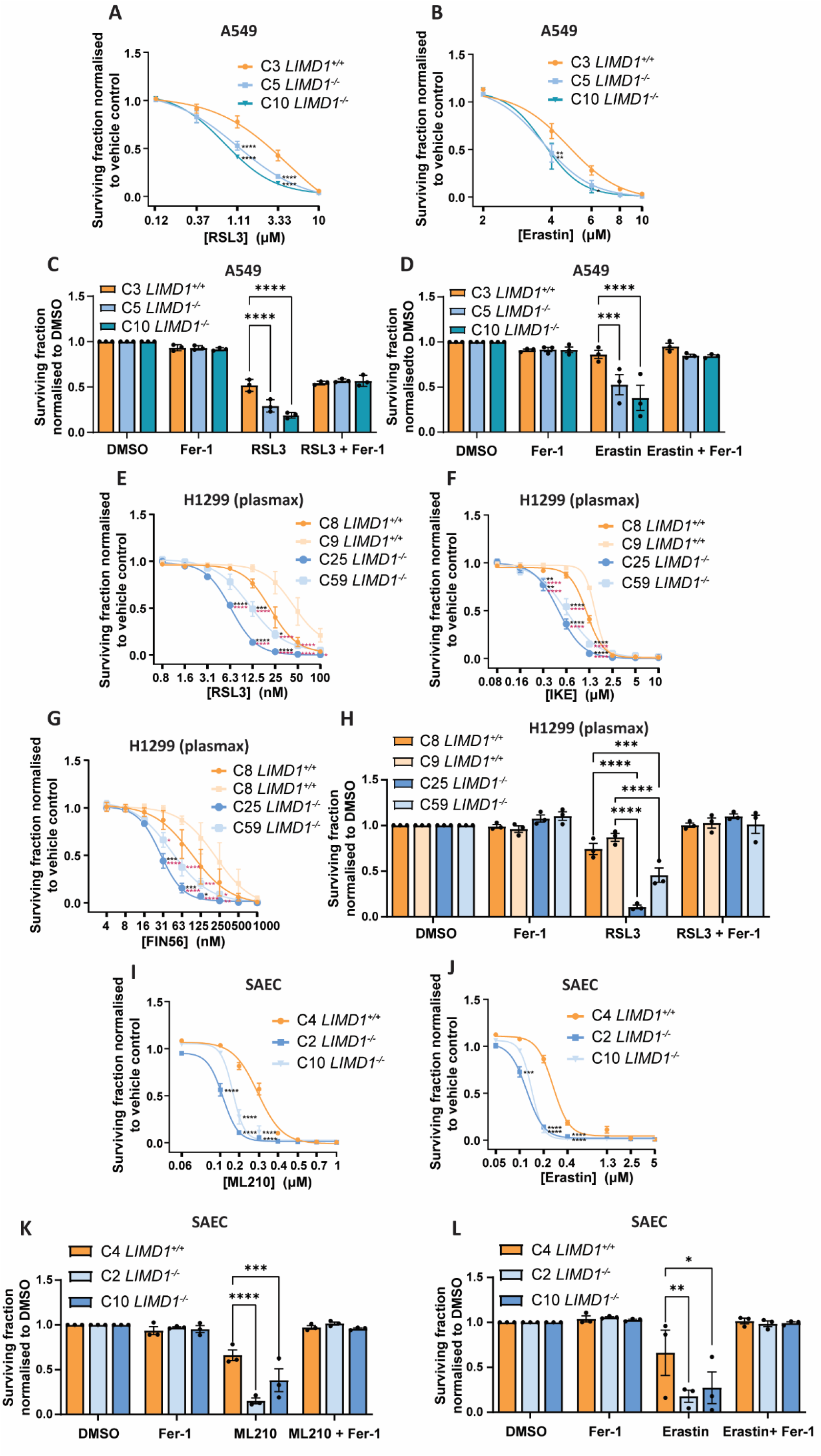
GPX4 targeting via ferroptosis inducing agents selectively inhibits *LIMD1*-deficient cells. **(A, B)** Dose-response curves showing surviving fractions of isogenic A549 cells upon RSL3 (n=4) **(A)** and erastin (n=2) treatment **(B)**, seeded at 1000 cells per well with a double dose of drug for 48 h followed by 48 h treatment. **(C)** Surviving fractions of isogenic A549 cells, seeded at 1000 cells per well, upon double dose of treatment of 48 followed by 48 h with DMSO, fer-1 (2 μM), RSL3 (2 μM), and RSL3 (2 μM) in combination with fer-1 (2 μM) (n=3). **(D)** Surviving fractions of isogenic A549 cells, seeded at 1000 cells per well, upon double dose of treatment of 48 followed by 48 h with DMSO, fer-1 (2 μM), erastin (5 μM), and erastin (5 μM) in combination with fer-1 (2 μM) (n=3). **(E-G)** Dose-response curves showing surviving fractions of isogenic H1299 cells grown in plasmax upon RSL3 **(E)**, IKE **(F)** and FIN56 **(G)** treatment for 24 h, seeded at 2000 cells per well (n=3). **(H)** Surviving fractions of isogenic H1299 grown in plasmax, upon 24 h treatment with DMSO, fer-1 (2 μM), RSL3 (8 nM) and RSL3 (8 nM) combined with fer-1(2 μM) (n=3). **(I, J)** Dose-response curves showing surviving fractions of isogenic SAEC cells upon ML210 **(I)** and erastin **(J)** treatment for 24 h, seeded at 3000 cells per well (n=3). **(K)** Surviving fractions of isogenic SAEC cells, upon 24 h treatment with DMSO, fer-1 (2 μM), ML210 (0.3 μM) and ML210 (0.3 μM) in combination with fer-1 (2 μM) (n=3). **(L)** Surviving fractions of isogenic SAEC cells, upon 24 h treatment with DMSO, fer-1 (2 μM), erastin (0.3 μM) and erastin (0.3 μM) in combination with fer-1 (2 μM) (n=3). Data is shown as biological replicates ± S.E.M. Two-way ANOVA with Dunnett’s multiple comparisons test was performed in **A-D, I-L**, whereas two-way ANOVA with Tukey’s comparisons test was performed in **E-H**, comparing mean values of C25 *LIMD1*^*-/-*^ and C59 *LIMD1*^*-/-*^ groups to both C8 *LIMD1*^*+/+*^, shown by black asterisk, and C9 *LIMD1*^*+/+*^ groups, shown by purple asterisk. \**p* ≤ 0.05, \*\**p* ≤ 0.01, \*\*\**p* ≤ 0.001, \*\*\*\**p* ≤ 0.0001.

As standard DMEM and RPMI lack sodium selenite, required for selenocysteine incorporation and full GPX4 activity ^37^, we validated our findings in Plasmax, a physiologically relevant medium containing 25 nM sodium selenite, ^38^. In H1299 isogenic cells cultured in Plasmax, *LIMD1*^-/-^ cells showed increased sensitivity to RSL3, IKE and FIN56 (Fig. 3E–G), all reversed by Fer-1 (Fig. 3H; Fig. S2A, B). Similar ferroptosis-specific sensitivity was observed in SAECs: ML210 and erastin selectively reduced the viability of *LIMD1*^-/-^ clones, with Fer-1 rescuing the effect (Fig. 3K–N). Together, these data robustly validate GPX4 as a ferroptosis-specific vulnerability in *LIMD1-*deficient lung epithelial cells across transformed and primary models.

### *LIMD1*-deficient cells exhibit a ferroptosis-prone basal state characterised by elevated lipid accumulation and oxidative stress

To understand the mechanisms underlying GPX4 dependence, we assessed key ferroptosis-enabling features. Phospholipid composition and oxidative stress are critical determinants of ferroptosis sensitivity: PUFA-rich membranes promote ferroptosis, while MUFAs are protective ^39-41^. Oil Red O staining revealed elevated basal lipid droplet accumulation in *LIMD1*^-/-^ H1299 cells, further exacerbated by BSA-oleate treatment (Fig. 4A, B), suggesting a differential lipid supply in *LIMD1* deficient cells.

**Figure 4.**
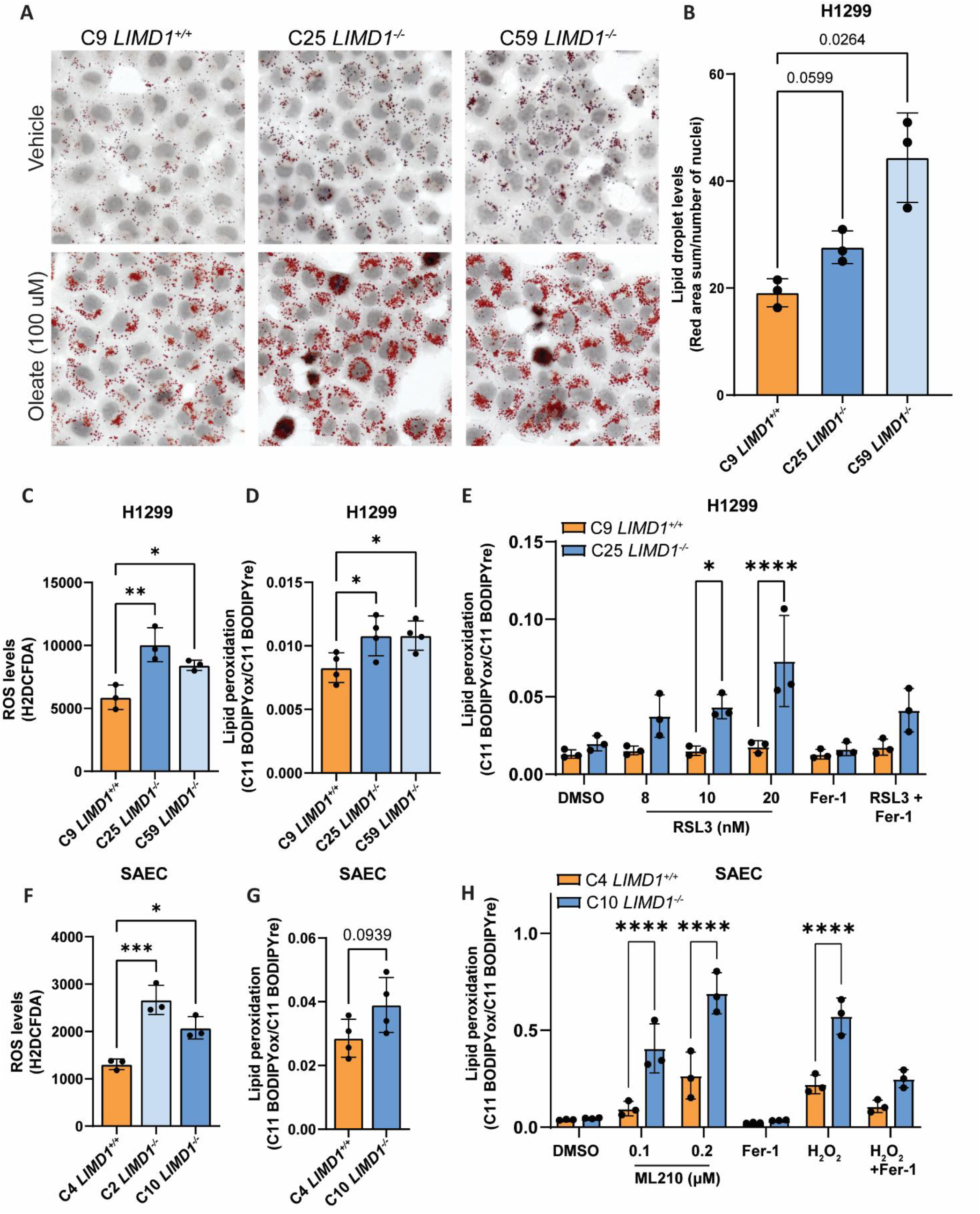
*LIMD1*-deficient cells exhibit higher levels of lipid droplets, ROS and lipid peroxidation. **(A)** Representative images of isogenic H1299 cells stained for lipid droplets with Oil Red O upon 24 h treatment with vehicle and BSA-conjugated Oleate (100 µM) (n=3). **(B)** Quantification of basal lipid droplet levels in isogenic H1299 cells. **(C)** ROS levels evaluated by H2DCFDA staining in isogenic H1299 cells following seeding in 6-well plates at a confluency of 1.25×10^6^ cells per well (n=3). **(D, E)** Lipid peroxidation levels evaluated by C11-BODIPY 581/591 staining in isogenic H1299 cells under basal conditions (n=4) **(D)** and upon treatment **(E)** with RSL3 (8 nM, 10 nM, 20 nM), fer-1 (2 μM) and RSL3 (20 nM) combined with fer-1 (2 μM) for 24 h (n=3). **(F)** ROS levels evaluated by H2DCFDA staining in isogenic SAEC cells following seeding in 6-well plates at a confluency of 1.25×10^6^ cells per well (n=3). **(G, H)** Lipid peroxidation levels evaluated by C11-BODIPY 581/591 staining in isogenic SAEC cells under basal conditions (n=4) **(G)** and upon treatment **(H)** with ML210 (0.1 µM, 0.2 µM), fer-1 (2 μM) for 24 h; H_2_O_2_ (300 μM) and H_2_O_2_ (300 μM) with fer-1 (2 μM) treatment was performed for the last 30 minutes (n=3). Data is shown as biological replicates ± S.D. in **C-H** and as biological replicates ± S.E.M. in **B**. One-way ANOVA with Dunnett’s multiple comparisons test was performed in **B-D, F**, whereas Two-way ANOVA with Bonferroni’s multiple comparisons test was performed in **E** and **H**. Two-tailed unpaired t-test was performed in **G**. \**p* ≤ 0.05, \*\**p* ≤ 0.01, \*\*\**p* ≤ 0.001, \*\*\*\**p* ≤ 0.0001.

We next assessed oxidative stress and lipid peroxidation using H2DCFDA and ratiometric C11 BODIPY (581/591) probes. *LIMD1*^-/-^ H1299 exhibited significantly elevated basal ROS and lipid peroxidation (Fig. 4C, D; Fig. S3A). Similar results were also obtained in the *LIMD1*^-/-^ SAECs, displaying significantly heightened basal ROS levels (Fig. 4F). RSL3 or ML210 treatment further increased lipid peroxidation in a dose-dependent manner only in *LIMD1*^-/-^ cells, again reversed by Fer-1 (Fig. 4E, H). Moreover, H_2_O_2_ exposure induced disproportionately higher lipid peroxidation in *LIMD1*^-/-^ cells, reinforcing their hypersensitivity (Fig. 4H). Reduced GSH:GSSG ratios in *LIMD1*^-/-^ cells confirmed elevated redox stress (Fig. S3B).

Collectively, these findings reveal a ferroptosis-prone state upon *LIMD1* loss, marked by increased lipid content, basal oxidative stress, and redox imbalance, providing mechanistic insight into the observed *GPX4* dependency.

## DISCUSSION

### GPX4 dependency reveals a synthetic vulnerability and therapeutic opportunity in *LIMD1*-deficient lung cancers

Lung cancer (LC) remains the most lethal cancer globally, with overall 5-year survival rates still dismal despite advanced in immunotherapy and targeted therapies. The urgent need for novel, biologically informed treatments in especially acute in non-small cell lung cancer (NSCLC), where tumour heterogeneity, late-stage diagnosis, and therapeutic resistance continue to undermine durable outcomes. Our study identifies and validates a critical vulnerability in *LIMD1*-deficient NSCLC–selective dependency on the selenoprotein glutathione peroxidase 4 (GPX4), offering a new therapeutic axis based on synthetic lethality.

*LIMD1* is frequently lost early in the pathogenesis on NSCLC, particularly in LUAD and LUSC subtypes. Loss of *LIMD1* occurs in a clonal fashion and correlates with reduced mRNA expression and poor patient prognosis ^16,17^. *LIMD1* deficiency dismantles multiple tumour suppressive circuits, but also presents a therapeutic challenge: no single downstream effector is easily druggable. Here, using unbiased CRISPR-Cas9 dropout screens in isogenic LUAD and primary airway epithelial cells, we uncover *GPX4* as a robust synthetic lethal partner in the context in *LIMD1* loss.

GPX4 plays a non-redundant role in detoxifying phospholipid hydroperoxides, safeguarding cells from ferroptosis ^26,28,31^. Our screens highlighted not only GPX4, but also components of the selenocysteine biosynthesis and incorporation pathway (EEFSEC, SEPSECS, PSTK), suggesting that *LIMD1*-deficient cells rely on a heightened ferroptosis defence program. We demonstrate that targeting GPX4 through siRNA or ferroptosis-inducing agents (FINs) such as RSL3, ML210, FIN56, and erastin, selectively kills *LIMD1*-deficient cells, both in LUAD lines and in primary SAECs. This synthetic lethality appears driven by a ferroptosis-sensitised basal state marked by elevated reactive oxygen species (ROS), enhanced lipid peroxidation, altered glutathione redox status, and increased lipid droplet accumulation.

This ferroptosis-prone metabolic rewiring has clear therapeutic implications. First, we show that *LIMD1*-deficient cells are selectively sensitive to three classes of FINs, underscoring the robustness of the *GPX4* dependency. Second, this sensitivity was retained in physiologically relevant media (Plasmax), supporting translational evidence. Third, rescue with the lipophilic antioxidant Fer-1 confirmed the observed cytotoxicity is ferroptosis-specific.

Importantly, our findings align with and extend previous reports linking GPX4 overexpression to poor outcomes in LUAD ^42,43^ and therapeutic resistance in NSCLC ^43-47^. In particular, GPX4 inhibition has been shown to synergise with cisplatin ^46^, reverse radiotherapy resistance ^47^, and enhance the effects of immune checkpoint blockade ^48^. This last point is especially notable: interferon-γ released by activated CD8^+^ T-cells downregulates SLC7A11 via STAT1, depleting glutathione and indirectly suppressing GPX4 function. Thus, combining FINs with immunotherapies in *LIMD1*-deficient tumours may offer a compelling dual-attack strategy that exploits both ferroptotic and immunogenic vulnerabilities.

From a biological perspective, our results point to a mechanistic model in which *LIMD1*-deficient cells accumulate oxidative and lipid stress, relying on GPX4 as a key survival mechanism. We demonstrate increased basal ROS and lipid peroxidation, as well as dysregulated lipid metabolism, both critical preconditions for ferroptosis ^39-41,49^. Notably, *LIMD1*-deficient cells also show enhanced lipid droplet formation and altered fatty acid composition, which could influence membrane vulnerability to peroxidation and ferroptotic commitment. These findings open the door to broader metabolic characterisation of *LIMD1*-deficient NSCLCs, including how PUFA uptake, incorporation, and organelle-specific redox homeostasis are rewired upon *LIMD1* loss.

### Future directions and translational outlook

To translate these findings into a therapeutic context, several next steps are essential. First, *in vivo* validation of *GPX4* dependency using FINs in LIMD1-deficient GEMM models and patient-derived xenografts will be critical to assess therapeutic efficacy and toxicity profiles. Second, lipidomic and metabolomic analyses will clarify which specific lipid classes and unsaturation profiles drive ferroptosis susceptibility in *LIMD1*-deficient cells. Third, high-resolution and organelle-specific imaging will help define where ROS and lipid peroxidation events initiate upon *LIMD1* loss and FIN challenge. Finally, understanding how LIMD1 interacts with the immune microenvironment may reveal further opportunities to combine ferroptosis inducers with immunotherapies or redox modulators.

## Conclusions

In sum, our study establishes GPX4 as a novel, actionable synthetic lethality in *LIMD1*-deficient NSCLC, revealing a ferroptosis-sensitive metabolic state that can be therapeutically exploited. By demonstrating selective GPX4 dependency across LUAD lines and primary epithelial models, and across multiple classes of FINs, we identify a tractable vulnerability with potential for personalised treatment strategies. As *LIMD1* loss occurs early and clonally in NSCLC evolution, targeting this axis may enable both early intervention and long-term disease control. These findings open new avenues for patient stratification and therapeutic design in one of the most challenging cancer types, and pave the way for clinical efforts to improve lung cancer outcomes through ferroptosis-based precision medicine.

## Acknowledgements

We acknowledge the technical assistance provided by the BCI Flow Cytometry and Microscopy Core Facilities. We thank the Cancer Research UK-AstraZeneca Functional Genomics Centre for conducting the CRISPR-Cas9 dropout screens.

---

**Supplementary Figure 1.**
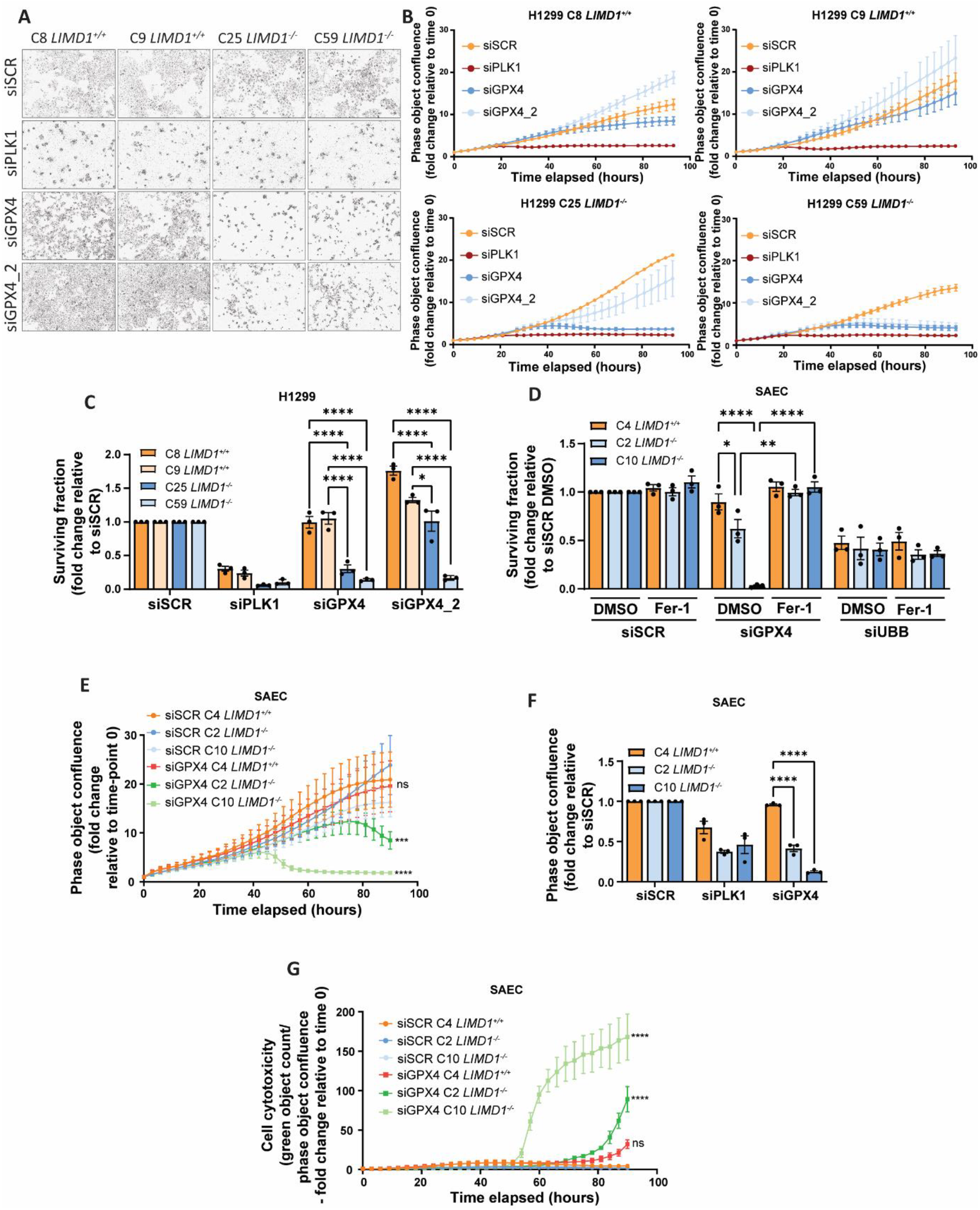
RNA interference-mediated *GPX4* targeting selectively inhibits *LIMD1*-deficient cells. **(A-B)** Representative images from live cell imaging of isogenic H1299 cells **(A)** and phase object confluence levels of isogenic H1299 cells **(B)**, seeded at 500 cells per well, upon treatment with 30 nM siRNA for 96 h (n=3). **(C)** Bar charts showing surviving fractions of isogenic H1299 cells, seeded at 500 cells per well, upon treatment with 30 nM siRNA for 96 h (n=3). **(D)** Surviving fraction of isogenic SAEC cells, seeded at 2000 cells per well, upon treatment with 30 nM siRNA for 96 h and treatment with DMSO and fer-1 (2 µM) 24 h post siRNA treatment (n=3). **(E)** Phase object confluence levels of isogenic SAEC cells, seeded at 2000 cells per well, upon treatment with 30 nM siRNA for 96 h. **(F)** Bar charts showing phase object confluence levels of isogenic SAEC cells, seeded at 2000 cells per well, upon treatment with 30 nM siRNA for 90 h (n=3). **(G)** Cell cytotoxicity levels of isogenic SAEC cells, seeded at 2000 cells per well, upon treatment with 30 nM siRNA for 117 h (n=3). Data is shown as biological replicates ± S.E.M. Two-way ANOVA with Tukey’s multiple comparisons test was performed in **C-E, G**. whereas two-way ANOVA with Dunnett’s multiple comparisons test was performed in **F**. \**p* ≤ 0.05, \*\**p* ≤ 0.01, \*\*\**p* ≤ 0.001, \*\*\*\**p* ≤ 0.0001.

**Supplementary Figure 2.**
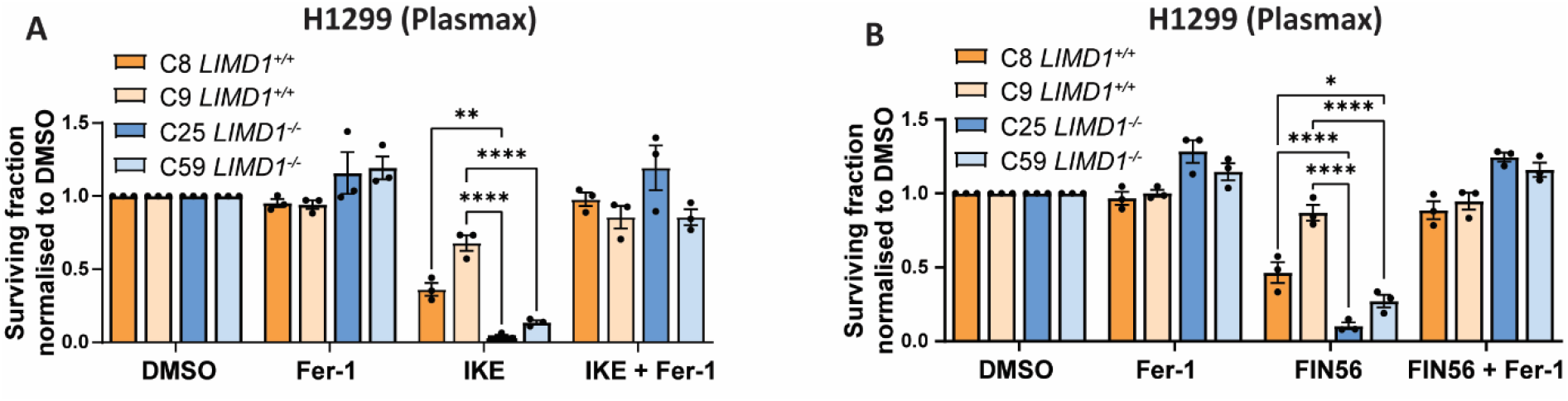
GPX4 targeting via ferroptosis inducing agents selectively inhibits *LIMD1*-deficient cells. **(A)** Surviving fractions of isogenic H1299 grown in plasmax, upon 24 h treatment with DMSO, fer-1 (2 μM), IKE (0.6 μM) and IKE (0.6 μM) combined with fer-1 (2 μM) (n=3). **(B)** Surviving fractions of isogenic H1299 grown in plasmax, upon 24 h treatment with DMSO, fer-1 (2 μM), FIN56 (0.06 μM) and FIN56 (0.06 μM) combined with fer-1 (2 μM) (n=3). Data is shown as biological replicates ± S.E.M. Two-way ANOVA with Tukey’s comparisons test was performed. \**p* ≤ 0.05, \*\**p* ≤ 0.01, \*\*\**p* ≤ 0.001, \*\*\*\**p* ≤ 0.0001.

**Supplementary Figure 3.**
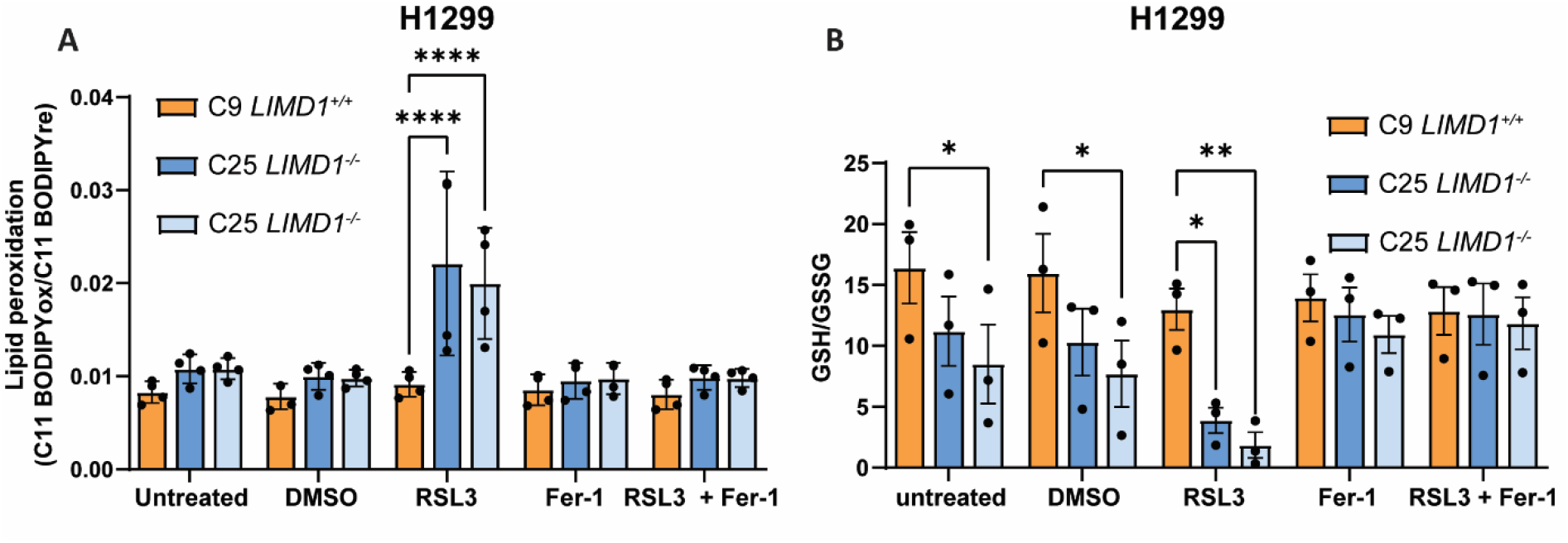
*LIMD1*-deficient cells exhibit higher degree of lipid peroxidation and oxidative stress. **(A)** Lipid peroxidation levels evaluated by C11-BODIPY 581/591 staining in isogenic H1299 cells upon treatment with RSL3 (20 nM), fer-1 (2 μM) and RSL3 (20 nM) combined with fer-1 (2 μM) for 24 h (n=4). Oxidative stress levels of isogenic H1299 cells, seeded at 2000 cells per well, upon treatment with RSL3 (10 nM), fer-1 (2 μM) and RSL3 (10 nM) combined with fer-1 (2 μM) for 24 h (n=3). Data is shown as biological replicates ± S.E.M. Two-way ANOVA with Dunnett’s comparisons test was performed. \**p* ≤ 0.05, \*\**p* ≤ 0.01, \*\*\**p* ≤ 0.001, \*\*\*\**p* ≤ 0.0001.

